# AniAnn’s: alignment-free annotation of tandem repeat arrays using fast average nucleotide identity estimates

**DOI:** 10.64898/2026.01.27.702063

**Authors:** Alexander Sweeten, Michael C. Schatz, Adam M. Phillippy

## Abstract

**Motivation:** Satellite DNA has long posed challenges for genome assembly and analysis due to its low sequence complexity and poor mappability. These large heterochromatic arrays of tandem repeats are ubiquitous across eukaryotic genomes, yet remain understudied. Current methods for annotating satellite regions, and other classes of tandem repeat arrays, are limited in their ability to annotate divergent or novel sequences.

**Results:** In this work, we introduce AniAnn’s, an algorithm for annotating large blocks of tandemly repeating DNAs. AniAnn’s exploits the high Average Nucleotide Identity (ANI) shared between repeat units of the same array to quickly and accurately infer the boundaries of such arrays. We show that AniAnn’s improves the annotation of satellites and other tandem repeats within a variety of plant and animal genomes, while requiring only a fraction of the runtime compared to previous approaches. We conclude by exploring several use cases of AniAnn’s as a lightweight method for masking repeats prior to whole-genome alignment as well as the *de novo* annotation and classification of satellite repeats.

**Availability:** AniAnn’s is open source software and available at github.com/marbl/anianns

## Introduction

As sequencing technology and assembly software continues to improve, completely gap-free “telomere to telomere” genomes have become the new standard in assembly quality. This is enabling the annotation of fully resolved satellite repeat arrays, which were mostly absent from prior eukaryotic reference genomes (Altemose et al., 2022; Yoo et al., 2025). For example, recent analyses of humans (Rhie et al., 2023) and other primates (Makova et al., 2024) have shown that satellite arrays can account for over 50% of their Y chromosome sequence. Annotating these regions in humans has proven extremely useful in understanding the function and evolution of centromeres (Logsdon et al., 2024), identifying novel satellite arrays (Hoyt et al., 2022), determining a mechanism of Robertsonian chromosome formation (de Lima et al., 2024), and better understanding regulation of ribosomal DNAs (rDNAs) (Franklin et al., 2025).

While advances in sequencing technologies and assembly software have enabled the high-quality reconstruction of satellite regions, accurately annotating them remains a difficult problem. Current state-of-the-art annotation methods, such as RepeatModeler and RepeatMasker (Flynn J et al., 2020; Smit et al., 2013-2015), require a manually curated library of repeats, such as Dfam or Repbase (Storer et al., 2021; Kojima et al., 2018) in addition to a local alignment scheme such as BLAST (Altschul et al., 1990). While these methods work well for well known repeats, they often fail to detect novel or highly diverged repetitive elements. In fact, satellite arrays are among the most variable regions of the genome, and frequently undergo mutation through unequal crossing over, replication slippage, and point mutation at rates higher than the rest of the genome (Rudd et al., 2006; Miga et al., 2021; Thakur et al., 2021).

In parallel, a variety of *a priori* satellite annotation methods are available to partially overcome the above problems. Tandem Repeats Finder (TRF) (Benson et al., 1999) uses a match/mismatch/indel scoring-based system to identify approximate repeats, but is limited to finding repeats less than 2,000 base pairs and is unable to detect Higher Order Repeat (HOR) structures. TRASH (Woldzmeirez et al., 2023) detects tandem repeats within highly repetitive windows of a genome by aligning to a generated consensus repeat motif using MAFFT (Katoh et al., 2002). However, this approach fails for divergent satellite arrays, and has difficulty annotating composite repeats such as human β and γ satellites. RepeatOBserver (Elphinstone et al., 2025) and Spectral Repeat Finder (Sharma et al., 2004) both exploit the inherent repetitiveness within satellite arrays by mapping the sequence into the frequency domain using the Fourier transform to infer their locations and periodicity. However, because Fourier-based methods rely on detecting consistent periodic signals, their sensitivity is reduced for highly divergent or composite arrays.

In our previous work, we developed the software ModDotPlot (Sweeten et al., 2024) to rapidly identify and visualize tandem repeats within an assembled genome. ModDotPlot estimates the Average Nucleotide Identity (ANI) between sets of *k*-mers, using a modimizer-based variant of the Mash Screen containment score (Ondov et al., 2019). Since satellite repeat units within an array tend to share high ANI, regions of high self-similarity identified by ModDotPlot can be inferred as satellite arrays, without the need for a repeat database. However, ModDotPlot provides only approximate genomic locations, can be slow to run with the sensitivity needed to detect short arrays, and is unable to classify the arrays by satellite type.

In this work, we describe AniAnn’s: a fully automated and alignment-free satellite array annotation pipeline based on the modimizer analysis scheme of ModDotPlot. AniAnn’s combines image processing and segmentation algorithms with *k*-mer based operations to accurately infer tandem repeat array boundaries. This approach is generalizable to tandem arrays of any repeating element, including rDNAs and transposable elements (TEs), so we refer here to all such regions as “satellites” for simplicity. By using pairwise ANI of genomic windows to identify tandem arrays, AniAnn’s bypasses the problematic steps of database search and alignment, which limit the sensitivity and speed of prior approaches. We show that this alignment-free approach results in substantially improved runtime and memory usage, and a greater ability to detect diverged arrays. To conclude, we demonstrate the application of AniAnn’s on several recently completed T2T genomes.

## Materials and Methods

AniAnn’s takes as input a list of sequences in FASTA format, and computes a self-similarity matrix of ANI values for each sequence. To produce this matrix, we replicate the approach described in our previous work ModDotPlot (Sweeten et al., 2024). Briefly, we partition an input sequence into windows that are then decomposed into sets of *k*-mers, and compute an estimate of the ANI between pairwise combinations of sets using a min-hash style estimate of the containment index (Broder, 1997, Ondov et al., 2019). Prior to computing the containment index, each set is sketched using the modimizer scheme, which includes *k*-mers in the set if they are exactly divisible by a sparsity parameter *s*. This probabilistic sampling via modimizers enables efficient scaling to very large sequences (>100 Mb). We note that AniAnn’s is packaged as its own independent Python package and does not require ModDotPlot as a software dependency.

After computing the ANI matrix, AniAnn’s infers the location of satellite arrays based on the formation of squares along the main diagonal of the matrix (Figure 1). Other rectangles that do not bisect the main diagonal, but do match to detected satellite array coordinates in either the *x* or *y*-axes, are used to link arrays by class. We perform further analysis to refine the boundaries of each satellite array, as well as determine their orientation and periodicity (i.e. unit size). Each annotated array can then be evaluated against a *k*-mer database of known satellite classes, enabling assignment of a class to each region.

**Figure 1.**
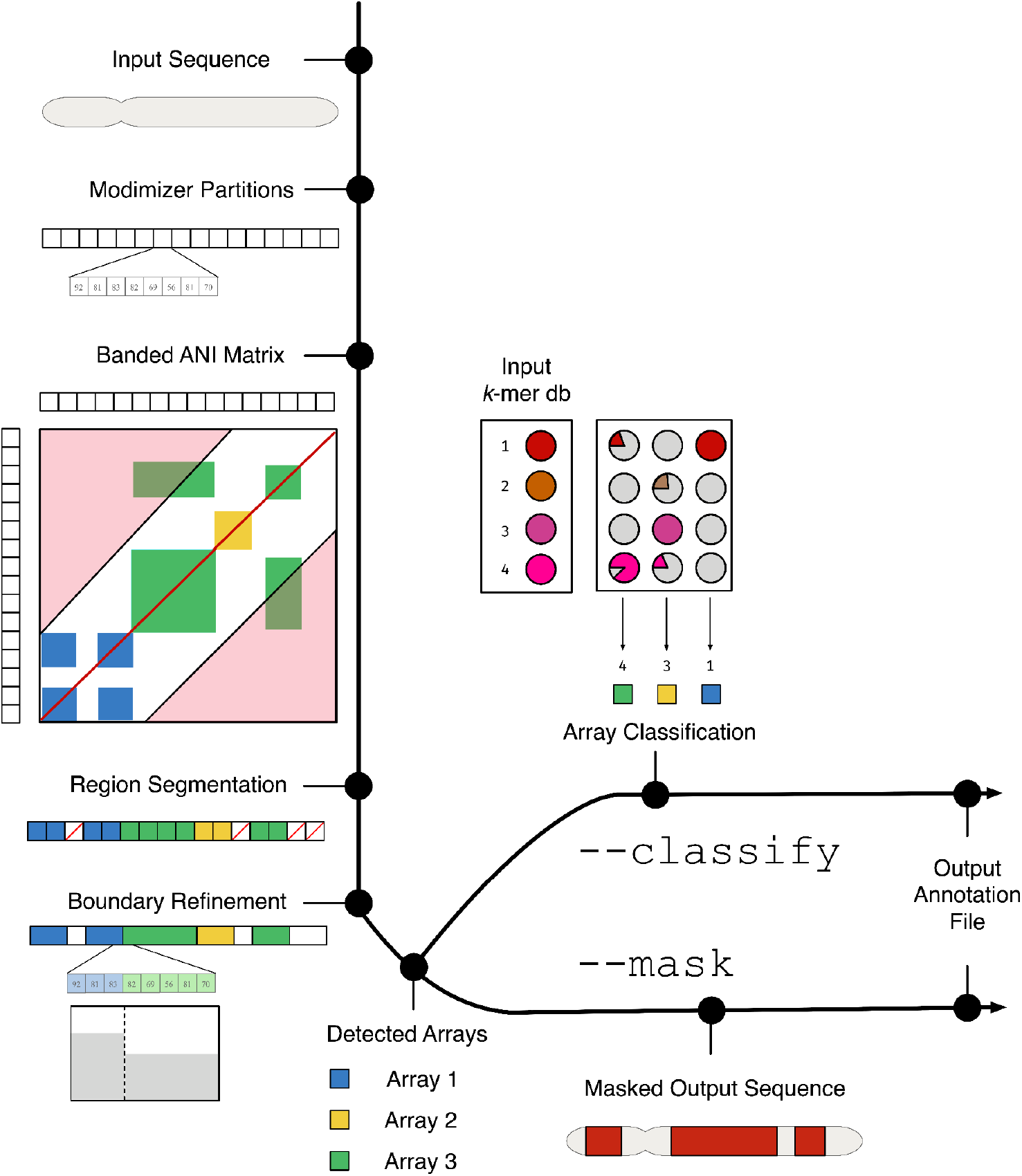
Overview of AniAnn’s pipeline. A sequence in FASTA format is provided as input, which is then partitioned into sets of modimizers. After conversion, a banded ANI matrix *M* is computed (regions in red are excluded from the band). The matrix *M* is used to infer regions containing satellites. The inferred regions are post-processed using *k*-mers to refine boundary points and link related arrays to each other. If desired, the user can provide a database of known satellite *k*-mers with the `--classify` parameter. If this option is selected, selected satellite arrays will be classified according to their highest match in the database. If the `--mask` parameter is selected, AniAnn’s will mask the input FASTA based on the regions detected in *M*.

### Diagonal Banding

Given a sequence of length *n* partitioned into windows of size *w*, we define the number of partitions *r* = (*n* − *k* + 1)/*w* (also called the resolution), where *k* is the *k*-mer length. Each partition includes approximately *w*/*s* sampled *k*-mers from which the pairwise ANI of any two partitions can be estimated. Computing the upper-triangle of a self-similarity matrix *M* requires 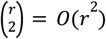 ANI calculations. Despite having a quadratic dependency on *r*, creating a self-similarity matrix can be fast in practice by keeping *w* sized proportionally to *n*, effectively keeping the plot resolution constant, e.g. both a 1 Mb or 100 Mb sequence are divided into *r* = 1000 partitions. This is effective for visualization purposes, as is the case for ModDotPlot; however, if *w* is significantly larger than the length of the array, then the array will be undetectable in the ANI matrix. The solution of decreasing *w* (thus increasing *r*) can lead to unacceptably slow runtimes on large sequences.

To address this complexity problem, observe that all satellite arrays longer than *w* appear as a high-identity square bisected by the *x* = *y* diagonal of a self dotplot. Thus, it is sufficient to only explore a banded region around the diagonal to identify such regions. Formally, given an *r*-by-*r* matrix *M* and a band limit *b*, we compute *M*[*i,j*] only if |*i* − *j*| ≤ *b*. Treating *b* as a tunable parameter allows a trade-off between sensitivity and runtime; when *b* is fixed to a constant independent of *r*, the runtime is reduced to *O*(*r*). Banding does not reduce the sensitivity of satellite detection, but it does remove information regarding the relatedness of similar satellites (i.e., as would be captured by the off-diagonal entries). However, as we show later, these similarities can be separately computed after the initial identification of the satellite regions.

### Region Segmentation

Given a self-identity matrix *M* (Figure 2a), AniAnn’s applies the Sobel-Feldman operator (Sobel et al., 1990), which convolutes *M* with weighted horizontal and vertical gradient filters *G*_*x*_ and *G*_*y*_. The output of this process is a binary edge map *M’* that clearly highlights detected edges, making inference of satellite boundaries easier to detect (Figure 2b). *M’* is further decomposed into two complementary components: *M’*_*diag*_ and *M’*_*distal*_. *M’*_*diag*_ contains only valid squares bisected by the *x* = *y* diagonal, representing satellite arrays. *M’*_*distal*_ contains the remaining rectangles that do not intersect the main diagonal, representing similarity between distant satellite blocks (Figure 2c).

**Figure 2.**
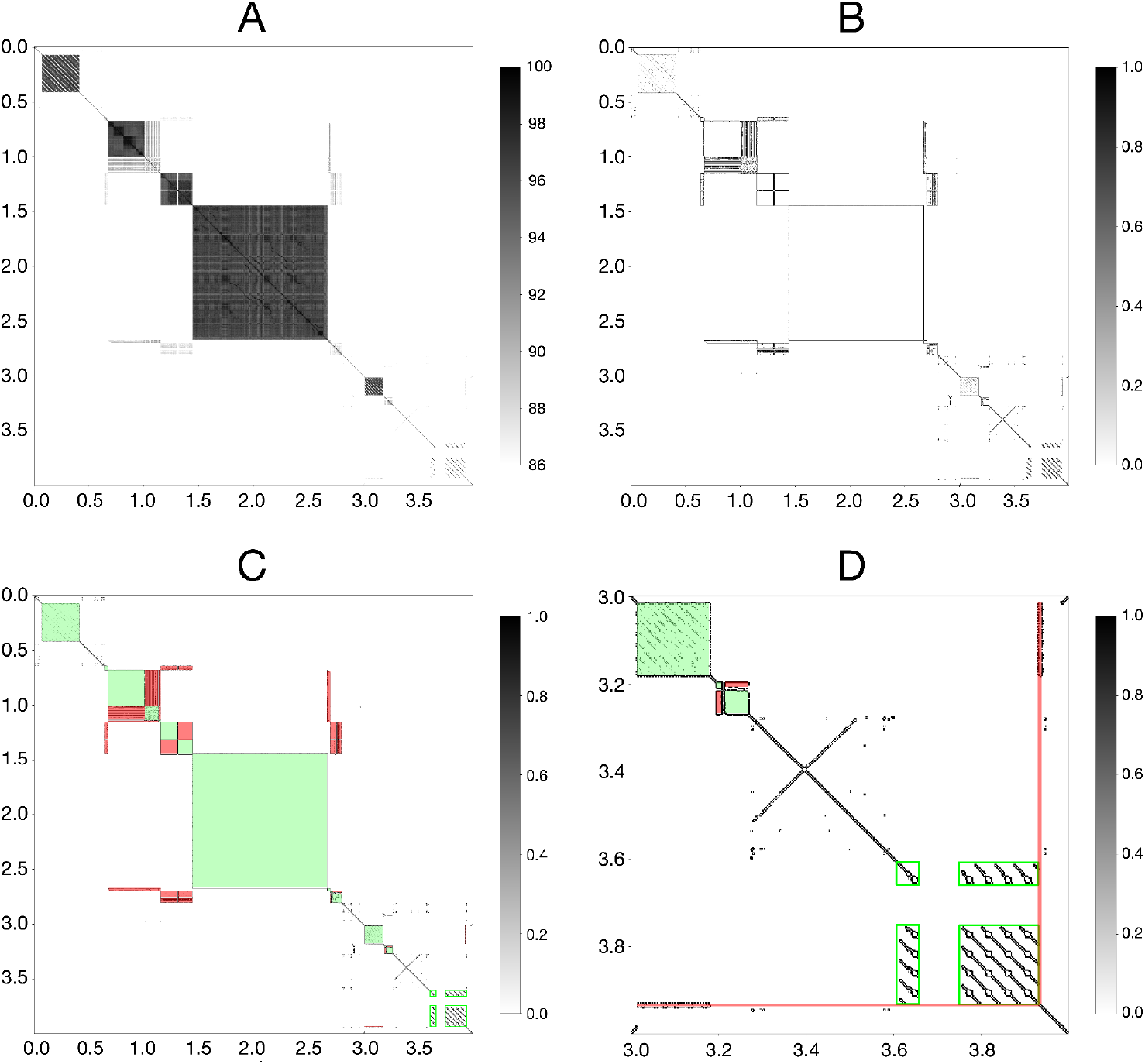
A) ANI matrix *M* of HG002 Chr13_MATERNAL:1–4000000, visualized as a grayscale heatmap with values ranging from 86 to 100. B) After applying the Sobel-Feldman operation to *M*, the output *M’* becomes a binary edge map, enabling easier edge detection. C) *M’* is further split into *M’*_*diag*_ (green) and *M’*_*distal*_ (red), representing satellite arrays and their homology relationships, respectively. D) A zoomed in plot of *M’* highlighting Chr13_MATERNAL:3000001–4000000. rDNA (light green boxes) is detected flanking a gap region between 3.6–3.9 Mb. Additionally, a single repeat copy not detected in *M’*_*diag*_ is inferred in *M’*_*distal*_ (red).

Any square in *M’*_*diag*_ is stored as a tuple (*x*_*0*_, *x*_*1*_), where *x*_*0*_ and *x*_*1*_ represent the start and end positions of each array, determined by their *x*-axis position in *M’*_*diag*_. Each tuple is added as a disjointed set into a Disjoint Set Union (DSU) data structure (Galler et al., 1964). Using *M’*_*distal*_, the bounding box of any existing rectangle is identified. If both the *x*-axis positions of the rectangle match with the start and end positions of any tuple in the DSU, and the *y*-axis positions of the rectangle match with the start and end positions of a different tuple, we apply the DSU union_set operation on the two matching sets. In the case where there is a match in one tuple but not any other (i.e., the *x*-axis positions match an element in the DSU but not the *y*-axis, or vice-versa), AniAnn’s adds a new tuple in the same set. We show an example of this occurring in Figure 2d, where a single copy of the ACRO repeat unit is inferred in *M’*_*distal*_, despite only matching to one position in *M’*_*diag*_. This case can occur when detecting very small arrays (i.e., substantially smaller than the window size), or when a single copy within the band of a larger, multi-copy array (Figure 2d).

If the length of a repeating unit within a satellite array is greater than the window size, then the assumption of a square in *M’*_*diag*_ is violated. Instead, lines parallel to the diagonal appear in *M’*_*distal*_ (Figure 2d). In our previous work (Sweeten et al., 2024), we discovered that inaccurate ANI estimates can arise between windows of size *w* that are “out of register” (i.e. have overhanging ends), so when estimating containment of one window within another, each containing window is expanded *w*/2 to the left and right to ensure the contained window will not have any overhang. Thus, the case of parallel lines will occur whenever the tandem unit size is greater than 2*w*. A good example of this is the human and primate rDNA arrays (Hori at al., 2021), which have a unit size of approximately 45 kb, which is substantially larger than *w*. To detect such tandems, we apply SciPy’s bounding box operation (Virtanen et al., 2020) specifically at a direction parallel to the diagonal. Collections of these bounding boxes intersecting with the diagonal are merged again in the DSU.

### Region Refinement and Orientation

Initially, each satellite array is assigned provisional boundaries based on its start and end coordinates (*x*_0_, *x*_1_), which occur at multiples of *w*. In practice, the true boundaries may shift anywhere within the ranges (*x*_0_ ± *w*/2, x_1_ ± *w*/2) due to expanding each window by a factor of *w*, as described above. Let *L* denote the *k*-mers falling within the possible left-boundary interval [*x*_0_ − *w*/2, *x*_0_ + *w*/2], and *R* denote the *k*-mers within the possible right-boundary interval [*x*_1_ − *w*/2, *x*_1_ + *w*/2]. We subdivide *L* and *R* into smaller windows (default size: *w*/20) and compute the containment index between the *k*-mers in each window and the “core” *k*-mers of the internal, non-border regions of the satellite array. We then identify the leftmost window in *L* and the rightmost window in *R* whose containment index exceeds a user-defined containment threshold (default: 0.2). The genomic position of the first *k*-mer in the selected *L* window is assigned as the left boundary, and the position of the last *k*-mer in the selected *R* window is assigned as the right boundary (Supplementary Figure 1). If every window in *L* or *R* exceeds the containment threshold, the DSU is queried to determine whether any neighboring arrays border *L* or *R*; if so, those arrays are merged and the boundary process is repeated on the merged array.

By default, canonical *k*-mers that consider the forward and reverse complement sequences together are used for both the creation of the ANI matrix and for adjusting detected array boundaries. If desired, the relative orientation of each array can be determined by replacing the “core” internal array *k*-mers described above with forward-strand *k*-mers only. To do this, we subdivide the core *k*-mers into windows (default size: *w*) and randomly select one candidate window to reverse-complement. The containment index is then computed between every core *k*-mer window and the candidate reverse complement window. Windows with containment values at or near zero indicate forward orientation (i.e. there are no reverse *k*-mers present in the candidate window), whereas windows with non-zero containment indicate reverse orientation. A suitable threshold for determining a forward versus reverse orientation is chosen using Otsu’s method (Otsu, 1979), which maximizes the variance between forward- and reverse-oriented windows (Supplementary Figure 2, Supplementary Figure 3).

### Satellite Classification

Inferred arrays are classified by comparing the *k*-mers within each set of the DSU to those from a reference database of known satellite *k*-mers. This database is generated by supplying AniAnn’s with an annotation file containing the genomic coordinates of established arrays, along with the corresponding sequence file. Because AniAnn’s uses a compact binary format and satellite arrays are highly compressible (few unique *k*-mers), the resulting *k*-mer representation takes up minimal space. As an example, the full catalog of satellite-associated *k*-mers from the human haploid CHM13 reference genome (*k* = 21) requires less than 32 MB of storage (Supplementary Table 1).

Within each DSU-defined satellite cluster, all *k*-mers from the constituent arrays are aggregated into a single query set. To reduce the size of large sets and keep each query relatively consistent, this set is also downsampled using modimizers with a default sketch size of 100,000 (Sweeten et al., 2024). The resulting modimizer sketch is compared against each reference satellite database using the containment index to measure how well the query *k*-mers are represented within each reference set. Each DSU cluster is then assigned to the satellite type corresponding to the database yielding the highest containment score passing a user-defined containment threshold (default: 70%). Clusters that fail to reach the containment threshold are conservatively left unclassified.

After completing the DSU of satellite array locations and refining their boundaries, AniAnn’s determines each array’s canonical repeat size. For this, we apply the NTRPrism algorithm (Altemose et al., 2022) to obtain a distribution of *k*-mer intervals. In brief, NTRPrism decomposes a sequence into *k*-mers using a deliberately small *k* value (default *k* = 6), which we refer to as *interval k-mers* (to differentiate them from the *k*-mers used for ANI computation and boundary refinement). For each interval *k*-mer that appears more than once within an array, the distance between its nearest exact match is recorded. As a result, in the histogram a peak occurs at the canonical repeat length *L*, while the height of peaks at higher harmonic intervals (2*L*, 3*L*, 4*L*, …) diminishes with *n* (Figure 3). In the case of a HOR, which consists of *n* nested tandem repeats, the *n*^th^ harmonic peak in the histogram will be disproportionately higher than its neighbors. For each harmonic peak *H*_*n*_, we compute a local enrichment ratio *R* _*n*_ = 2*H* _*n*−1_/(*H* _*n*_ + *H* _*n*+1_). For any *n* > 1, if *R*_*n*_ ≥ 2, we infer that the satellite array is organized as a higher-order repeat (HOR) and annotate it with the corresponding base repeat length *L* and HOR multiplicity *n* in the output annotation file.

**Figure 3.**
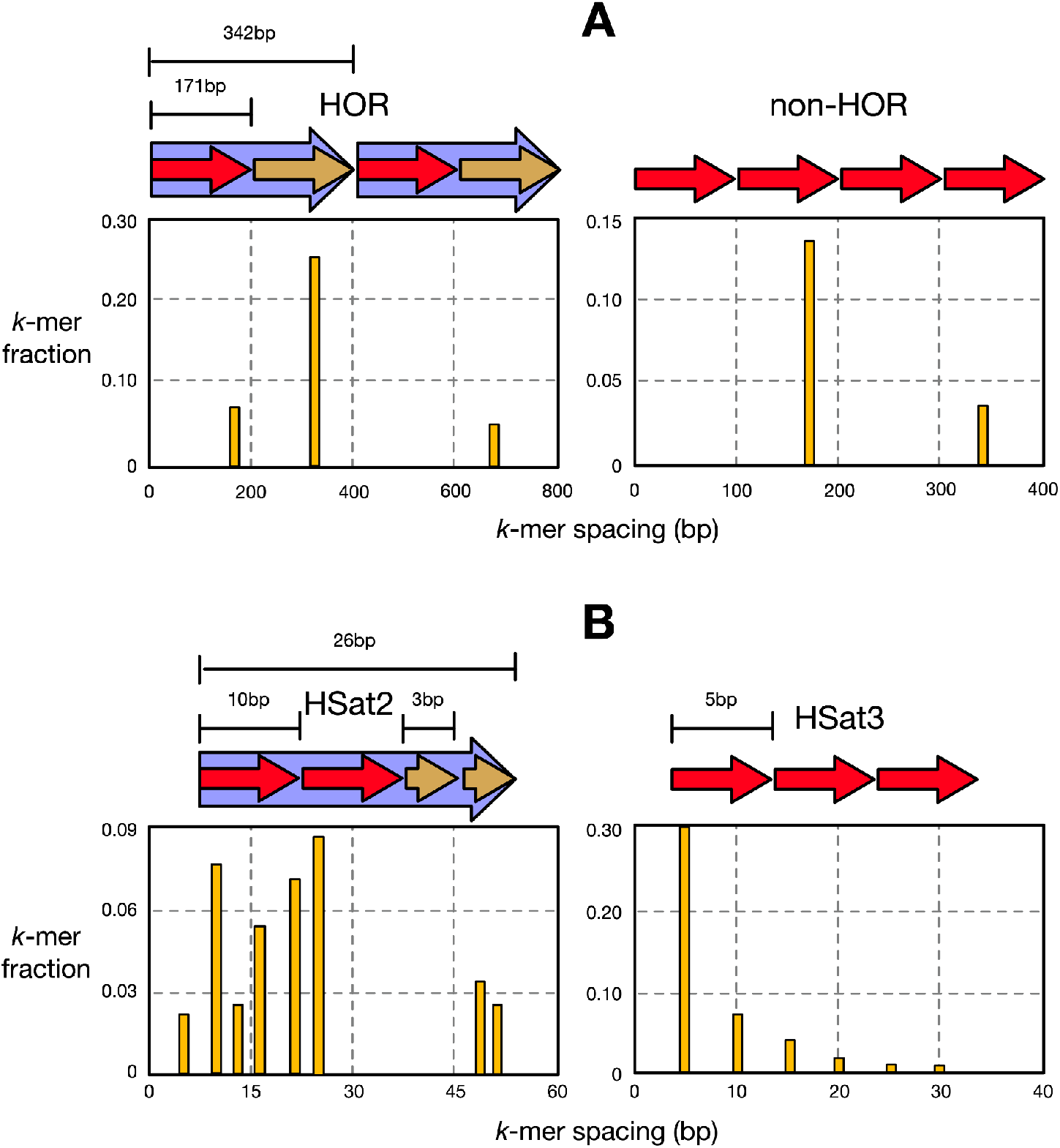
Visualization of NTRPrism on various human αSat and HSat arrays. (A) Histograms of spacings between *k*-mers within αSat arrays. In a HOR (left), peaks form at intervals based on the length of the αSat monomer (171 bp). The largest peak corresponds to the length of the HOR, occurring at a multiple of the monomer length (171 ^*^ 2 = 342 bp). When not arranged as a HOR (right), the tallest peak simply occurs at the monomer length and the peaks diminish with higher multiples. (B) Histograms of spacings between hSat2 (left) and hSat3 (right) arrays. HSat2’s canonical template consists of two identical decamers preceding one to two identical trimers, resulting in nested peak heights. This contrasts with HSat3 consisting of a single pentamer.

In cases where a query exhibits comparable containment across multiple satellite databases, rendering a highest-score assignment ambiguous, we use interval *k*-mers as described above to resolve the classification. Such ambiguity typically arises among closely related satellite families whose *k*-mer compositions are sufficiently similar to promote DSU merging. For example, human HSat2 and HSat3 both originate from the simple (CATTC)_*n*_ motif and can be difficult to distinguish. When finer-grained discrimination is required, we compute the cumulative distribution function (CDF) of interval *k*-mer distances for the query and for each top-matching reference database. A two-sample Kolmogorov–Smirnov test is used to identify the satellite class whose interval-length distribution most closely matches that of the query (Supplementary Figure 4).

## Results

### Satellite Classification Accuracy

To evaluate the accuracy of identifying and classifying satellite arrays, we apply AniAnn’s on several recently assembled genomes with high-quality annotations including *Homo sapiens* (HG002), *Gorilla gorilla* (mGorGor1), and *Luzula sylvatica* (IpLuzLuz). We benchmarked the results from AniAnn’s against TRASH2 (https://github.com/vlothec/TRASH_2), another method for *de novo* satellite identification. Both tools were first run in *de novo* mode, i.e., without using a satellite *k*-mer database (in the case of AniAnn’s) or template satellite sequences (TRASH2). Each tool was then run utilizing their respective classification method, with final accuracy evaluated using the intersection of the two result sets. This intersection was chosen to reduce false positives and ensure that only consistently detected arrays were retained for evaluation. Data locations and command line arguments for these experiments are available in Supplementary Tables 1 and 2 respectively.

We first examined the five most common satellite classes present on the complete human diploid genome HG002 (Figure 4a), with each class making up tens of megabases of sequence. Across both tools, classification metrics were greatest in low-divergence satellite arrays and tended to worsen in arrays with higher levels of sequence divergence. This trend of performance decline in higher-divergence arrays was more pronounced in TRASH than in AniAnn’s. Importantly, HSat2 and HSat3 are differentiated with a high rate of accuracy in both tools, although AniAnn’s shows overall higher performance by up to 5% gains in F1. This is notable as tools such as RepeatMasker have previously struggled to accurately classify these arrays, as they derive from the same ancestral motif (Altemose, 2022).

**Figure 4.**
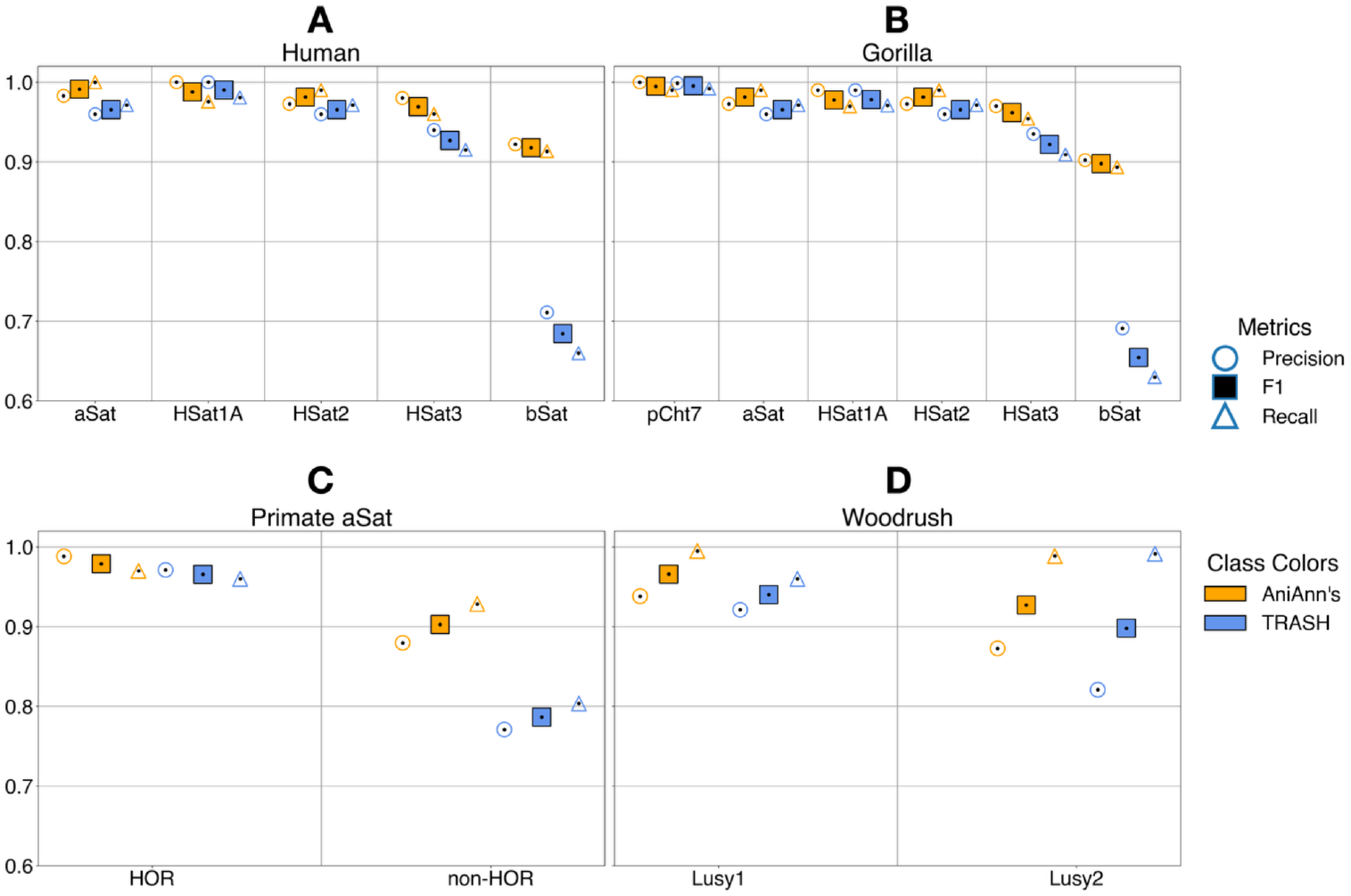
Classification accuracy metrics of both AniAnn’s and TRASH, across various organisms and satellite classes. A) Main human satellite classes using the diploid HG002 reference genome assembly. B) Main gorilla satellite classes using the diploid mGorGor1 genome assembly. C) Classification of αSat into HOR or non-HOR forming arrays, using the mGorGor1 and HG002 diploid assemblies. D) Main *Luzula sylvatica* (woodrush) satellite classes using the haploid lpLuzSylv assembly.

Moreover, while AniAnn’s displayed improved accuracy across all array types, it substantially improved over TRASH2 in classifying βSat by over 25% gains in F1 (Figure 4a, 4b). In addition to displaying both high intra-array and inter-array sequence divergence (Supplementary Figure 5), βSat commonly forms a tandemly repeating composite structure with the LSAU satellite and various subunits (Hoyt et al., 2022). Investigating this further, we found that TRASH2 was systematically unable to detect the D4Z4 array, a large ∼3.2 kb βSat composite containing the ampliconic *DUX4* gene located in the subtelomeric q-arms of chromosomes 4 and 10. These large arrays are classified as βSat and were accurately detected by AniAnn’s. TRASH2’s internal selection criterion ignores windows failing to meet a required repetitiveness ratio, which is apparently problematic for larger repeat units such as D4Z4.

In our analysis of the major satellite classes of the complete gorilla genome, we included the subterminal heterochromatic caps. These regions, specific to the *Gorilla* and *Pan* lineages, are dominated by large arrays of the pCht7 repeat (32 b), and make up ∼13% of the Gorilla genome (Yoo et al. 2025), corresponding to over 425 Mb of sequence. Both annotation methods achieved >99% F_1_ score in classifying this repeat. Classification metrics for the remaining satellite classes were similar to those observed in HG002, with AniAnn’s consistently outperforming TRASH across classes, and both tools showing reduced performance with increasing intra- and inter-array sequence divergence.

The centromeres of human and the other great apes comprise an “active” αSat HOR array that interacts with the kinetochore, and is typically flanked by diverged or monomeric αSat copies (Altemose et al., 2022). While most primate HORs consist of αSat, non-αSat arrays can also occasionally form HOR structures (Altemose, 2022). As a final classification measure, we ran AniAnn’s and TRASH on the centromeres of human and gorilla to classify the identified αSat arrays as HOR or non-HOR (Figure 4c). In HOR regions, we further classified arrays by superfamily αSat (SF) and computed their periodicity (Supplementary Table 3, Supplementary Table 4).

Of the ∼378 Mb of αSat sequence present across 98 chromosomes, over 372 Mb is detected across both tools, with ∼15% of αSat sequence not forming a HOR, consistent with the proportions reported through extensive manual curation in (Yoo et al. 2025) (gorilla) and (Altemose et al., 2022) (human). Both tools achieved similar classification accuracy in detecting HOR forming αSat, with a bias towards undercalling HOR regions, as indicated by higher precision relative to recall. TRASH employs a similar algorithm to NTRPrism for its own HOR detection, with both tools showing 100% concordance in reporting the periodicity and SF of each HOR-bearing array (Supplementary Table 3, Supplementary Table 4). Detection of non-HOR–forming αSat was less accurate across both tools, reflecting its higher inter- and intra-array divergence (Supplementary Figure 5).

We further extended our analysis to the holocentric plant *Luzula sylvatica*, whose genome is composed of more than 35% satellite arrays. In contrast to primate genomes, where satellites are typically concentrated within a single centromeric region, *L. sylvatica* exhibits centromeric activity distributed along the entire length of each chromosome, allowing us to evaluate satellite detection performance in a genome with dispersed satellite organization. Two dominant satellite families: Lusy1 (124 b) and Lusy2 (174–175 b), form the primary components of the centromeric CENH3-binding regions (Mata-Sucre et al., 2024). AniAnn’s improves both precision and recall in detecting Lusy1 and Lusy2 (Figure 4d). Notably, although Lusy2 exhibits substantially higher inter-array sequence divergence than Lusy1 (Supplementary Figure 6), AniAnn’s reduces false positives, achieving over a 5% increase in precision relative to TRASH. Together, these results indicate that sequence divergence alone does not limit AniAnn’s ability to detect and classify centromeric satellite arrays.

### Repeat Masking

A common use case of repeat detection software is to mask repeats for other downstream applications, such as sequence alignment and read mapping. Tandem repeats can increase runtime and reduce alignment quality because their repetitive structure creates many nearly identical alignment targets across the sequence. For this reason, both BLAST and minimap2 pre-mask low-complexity regions using the DUST algorithm (Morgulis et al., 2006), with minimap2 additionally preventing high-copy *k*-mers from being selected as alignment anchors (Li, 2018).

To evaluate the effectiveness of AniAnn’s as a repeat masking tool, we benchmarked against other repeat masking software, including TRF (Benson et al., 1999), TRASH2, and longdust (Li, et al., 2025). To evaluate each tool, we masked HG002’s Chr21 maternal and paternal haplotype, and then aligned the result using lastz (Harris, 2007). Human Chr21 is an acrocentric chromosome with a strong enrichment of satellite repeats on its short arm. Figure 5 illustrates the total number of bases masked by each tool, compared against the reference satellite annotation of HG002. We show that AniAnn’s masks a similar percentage of sequence as other tools, while maximizing the overlap with known centromeric satellite (CenSat) regions. We further report the runtime and memory requirements, along with masking accuracy metrics based on the CenSat annotation track in Table 1. Complete command line parameters for each tool are given in Supplementary Table 2.

**Table 1.** Analysis runtime, memory and accuracy results. All masking metrics are summed up across both haplotypes. Masking accuracy metrics are based on the CenSat annotation track as the ground truth. Masking percentage, precision, recall, and F1 score calculations are summed up across both haplotypes. Total aligned bases are computed from the resulting lastz MAF alignment file.

| Tool | Masking<br>CPU<br>time<br>(mm:ss) | Masking<br>% | Masking<br>Precision<br>% | Masking<br>Recall<br>% | Masking<br>F1 % | Alignment<br>CPU time<br>(mm:ss) | Total<br>CPU time<br>(mm:ss) | Total<br>Pipeline<br>Memory<br>(G) | Total<br>Aligned<br>Bases |
| --- | --- | --- | --- | --- | --- | --- | --- | --- | --- |
| Unmasked | N/A | N/A | N/A | N/A | N/A | 08:36 | 08:36 | **1.32** | **44,955,911** |
| AniAnn's<br>(default) | 03:12 | 13.51 | **96.84** | 96.80 | **96.82** | 02:19 | 05:31 | 2.11 | 44,948,111 |
| AniAnn's<br>(--identity<br>92) | 02:43 | 9.36 | 69.21 | **99.83** | 81.75 | 03:16 | 05:59 | 2.08 | 44,950,947 |
| longdust<br>(default) | **01:05** | 14.11 | 90.14 | 86.25 | 88.15 | 01:59 | **03:04** | 1.51 | 42,097,152 |
| longdust<br>(--score<br>0.3) | 03:24 | 16.08 | 93.76 | 78.76 | 85.61 | **01:45** | 05:09 | 1.69 | 42,043,109 |
| TRF<br>(default) | 14:51 | 14.25 | 89.28 | 84.57 | 86.86 | 02:08 | 16:59 | 1.82 | 42,099,152 |
| TRF<br>(-m 2000) | 26:12 | 15.21 | 90.31 | 78.91 | 85.16 | 02:21 | 28:33 | 1.82 | 42,027,085 |
| TRASH 2<br>(default) | 45:31 | 20.24 | 93.41 | 62.32 | 74.76 | 01:55 | 47:26 | 2.83 | 35,660,254 |
| TRASH2<br>(-m 2000) | 46:09 | **22.82** | 96.80 | 57.27 | 71.96 | 01:51 | 48:00 | 2.89 | 35,212,301 |

**Figure 5.**
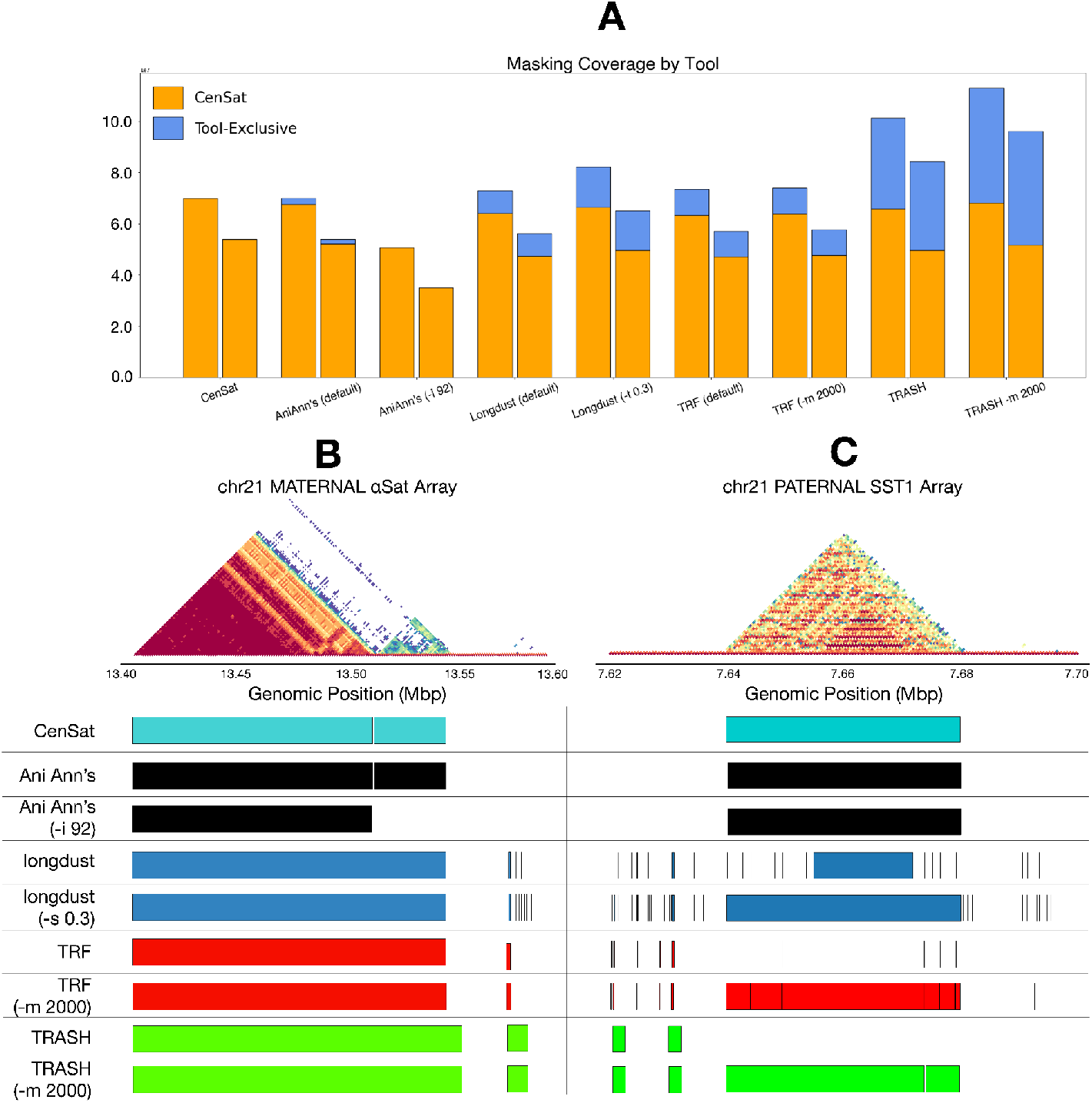
A) Number of bases of HG002 Chr21 masked by each tool, per haplotype. Percentage overlapping with the HG002 CenSat track is in orange, while remaining exclusive to each tool is in blue. ModDotPlot visualization of the maternal centromere, contained within chr21_MATERNAL:13400000–13600000. While all tools correctly mask αSat, only AniAnn’s is able to differentiate the HOR-forming array from the divergent non-HOR forming array without running the tool in a separate HOR-detecting mode. C) ModDotPlot visualization of the paternal SST1 array, contained within chr21_PATERNAL:7620000–7700000. AniAnn’s was the only tool able to completely mask the full SST1 array due to the longer unit size. Other tools are able to span this region only when modified from their default settings, which negatively impacts runtime and F1 score (Table 1).

From the T2T centromere satellite annotation, 14.77% of HG002’s Chr21’s maternal haplotype and 12.15% of its paternal haplotype are annotated as centromeric satellite regions. With the exception of TRASH2 which was over-masked by several megabases, similar percentages of sequences were masked from each remaining tool (Figure 5A). Under default parameters, AniAnn’s displayed the highest overlap with CenSat, maximizing the proportion of masked bases covered by the CenSat annotation while masking a similar amount of non-CenSat sequences as the other tools. Investigating these results further, we discovered a ∼47 kb array (chr21_PATERNAL:7638779–7681516) and a ∼103kb array (chr21_MATERNAL:11281392-11384751) of the SST1 satellite that were only partially masked by longdust, and completely missed by TRASH2 and TRF (Figure 5b). Both tools were able to detect this region upon increasing the default maximum repeat size to 2,000, but this caused the TRF runtime to double. The full SST1 array in longdust was detected by lowering the default complexity score threshold from 0.6 to 0.3, also at the expense of runtime.

Under default parameters, gapped extension of the pairwise alignment with lastz took over 48 CPU hours and required over 5 GB of memory when aligning the unmasked haplotypes, and over 4 hours after masking. To allow for the efficient testing of multiple tools and parameters, we therefore restricted our testing to only ungapped alignments (Table 1). Masking both haplotypes prior to alignment with lastz uniformly resulted in a ∼3–5x reduction in runtime across all masking methods with similar memory requirements. Thus, runtime of the masking method appears the most significant consideration for masking repeats prior to alignment, with default longdust parameters performing the best.

For our final analysis, we analyzed a three copy duplication on the paternal haplotype of HG002 Chr21 that is contained within an interspersed satellite region containing HSat3 and divergent αSat arrays (Figure 6). Under default parameters, all tools, including AniAnn’s, masked this entire region. As a result, when aligning masked versions of the haplotypes, information regarding structural variation is lost. ModDotPlot shows a high level of divergence between different regions of this array, ∼86–90% ANI, in contrast to >98% ANI for the SST1 example in Figure 5. As a result of this high intra-array divergence, a clear diagonal is visible in the dotplot between the two haplotypes, suggesting that they can be confidently aligned through this satellite region. Increasing the identity threshold of AniAnn’s to 92% unmasks this region, while keeping SST1 fully masked. This slightly increases alignment time compared to the default 86% identity threshold, but this tradeoff allows for the duplication variant to be captured by the alignment. Thus, because AniAnn’s provides ANI estimates for all identified satellites, it is possible in some cases to filter out the older homologies and isolate the orthology relationship. This provides an alternate means for identifying structural variants within satellite arrays, which is a notoriously difficult alignment problem (Bzikadze et al., 2023).

**Figure 6.**
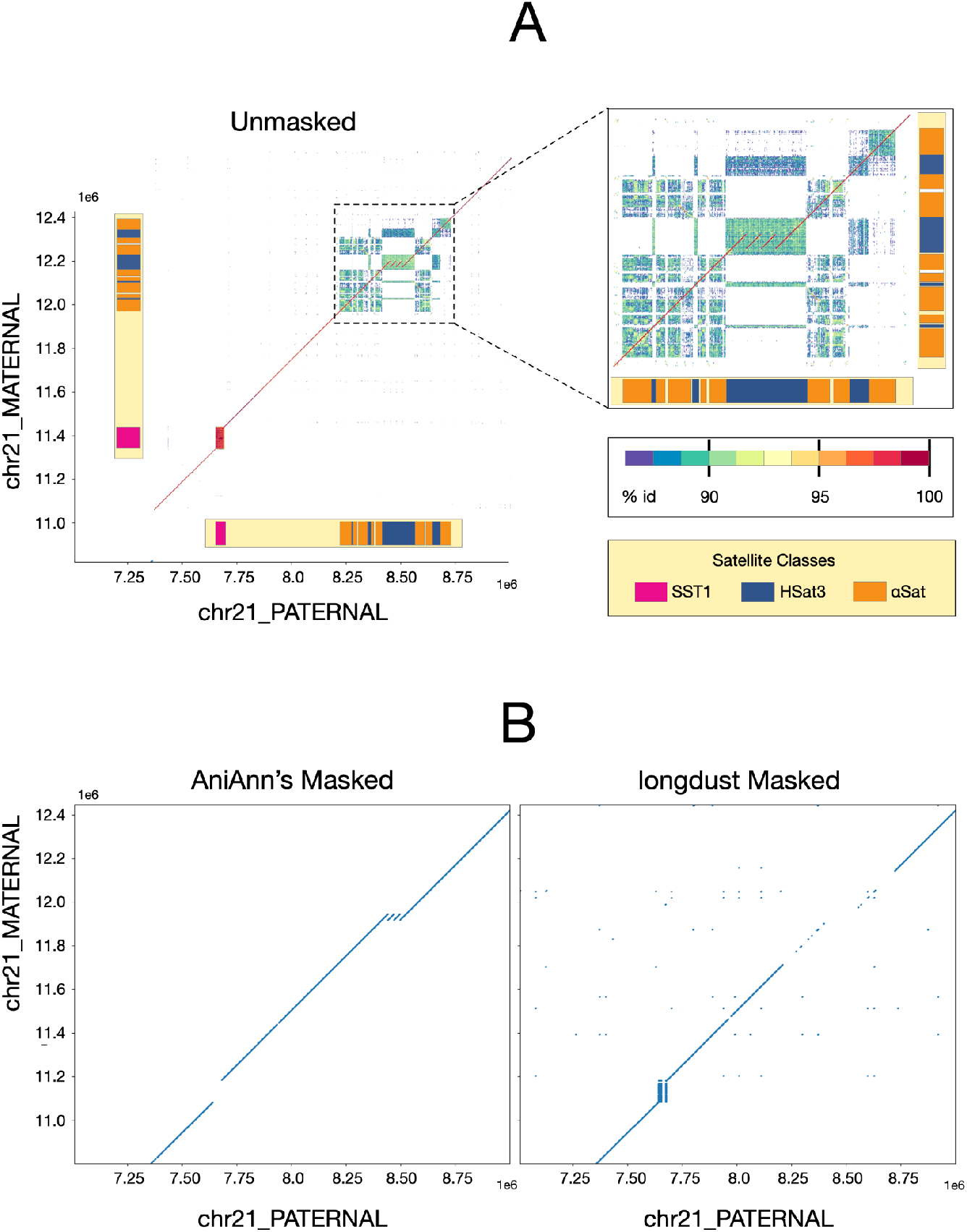
A) ModDotPlot comparative visualization of HG002 chr21_PATERNAL:7000000-9000000 and chr21_MATERNAL:11000000-12600000, annotated with various satellite classes. The zoomed-in plot highlights a three copy insertion embedded within an HSat3 array. ModDotPlot identifies the SST1 array as highly conserved (>98% idy), while hSat3 and αSat are significantly more diverged (<92% idy). B) Visualizations of lastz alignment of these regions masked with AniAnn’s under the `--identity 92` parameter (left) and longdust under default parameters (right). By not masking satellite arrays under 92% idy, alignment with the sequences masked by AniAnn’s is able to detect the highlighted insertion, whilst also masking the SST1 array.

## Discussion

Advances in long-read sequencing technologies have led to a rapid increase in the number of high-quality genome assemblies across a wide range of species, enabling the analyses of repetitive genomic regions that were previously inaccessible. This expansion has, in turn, highlighted the need for scalable and robust annotation methods capable of accurately characterizing complex repeat architectures in complete genomes. Standard alignment-based tandem repeat identification software is difficult to tune, has high runtime and memory requirements, and struggles to detect divergent or composite repeats. By focusing our attention on pairs of genomic windows that pass a minimum ANI threshold, AniAnn’s bypasses costly alignment steps while maintaining excellent satellite detection accuracy.

Our comparative analyses show that AniAnn’s consistently achieves high classification accuracy across diverse satellite arrays and genome architectures. Furthermore, by selecting for sequence identity as a parameter, AniAnn’s allows for greater control over the sensitivity of repeat masking, while remaining competitive with longdust in terms of speed. In human and gorilla, AniAnn’s exposes the full extent and organization of βSat and non-HOR αSat arrays, demonstrating that sequence divergence rather than array size or genomic context is the primary driver of missed annotations in traditional methods. These results show that accurate, genome-wide characterization of complex satellite architectures is now tractable without extensive manual curation.

We note that AniAnn’s does not currently attempt to identify a consensus repeat unit within satellite arrays. This step can be important for identifying internal array copy number variation and divergence. Nevertheless, we show that measuring repeat frequency using NTRPrism is sufficient in the detection of higher order repeats and differentiating related satellite classes. Given the speed of the identification and classification steps, AniAnn’s could be run as a prefilter before the construction of consensus sequences using custom methods such as HORmon [Kunyavskaya et al., 2020. PMID: 35545449].

While we focused here on satellite arrays, AniAnn’s has the potential to detect other types of repeats, including non-tandem repeats. As we have shown in our previous work, ModDotPlot reveals segmental duplications and palindromes as parallel and perpendicular off-diagonal lines respectively. Such structures are known to play important roles in genome function and evolution. Future AniAnn’s additions to identify and classify these distinct alignment patterns using related image analyses could provide an alignment-free approach for the quick annotation of segmental duplications throughout the genome, further enabling the analysis of genomic repeats across an ever-growing collection of complete genomes.

## Acknowledgements

We would like to thank Dmitry Antipov for helpful discussions regarding the methods sections of this paper, Hiram Clawson for suggesting improvements to the AniAnn’s codebase, and Hailey Loucks for providing the Centromeric Satellite Annotation for HG002 and mGorGor1. This work was supported, in part, by the Intramural Research Program of the National Human Genome Research Institute, US National Institutes of Health [to A.P.S. and A.M.P.]; NSF awards [IOS-2216612; DBI-2419522 to M.C.S.]. The contributions of NIH authors are considered Works of the United States Government. The findings and conclusions presented in this paper are those of the authors and do not necessarily reflect the views of the NIH or the U.S. Department of Health and Human Services. This work utilized the computational resources of the NIH HPC Biowulf cluster (https://hpc.nih.gov).

## Notes

### Competing Interest Statement

The authors have declared no competing interest.

