## Supplementary Figures for "AniAnn’s: alignment-free annotation of tandem repeat arrays using fast average nucleotide identity estimates"

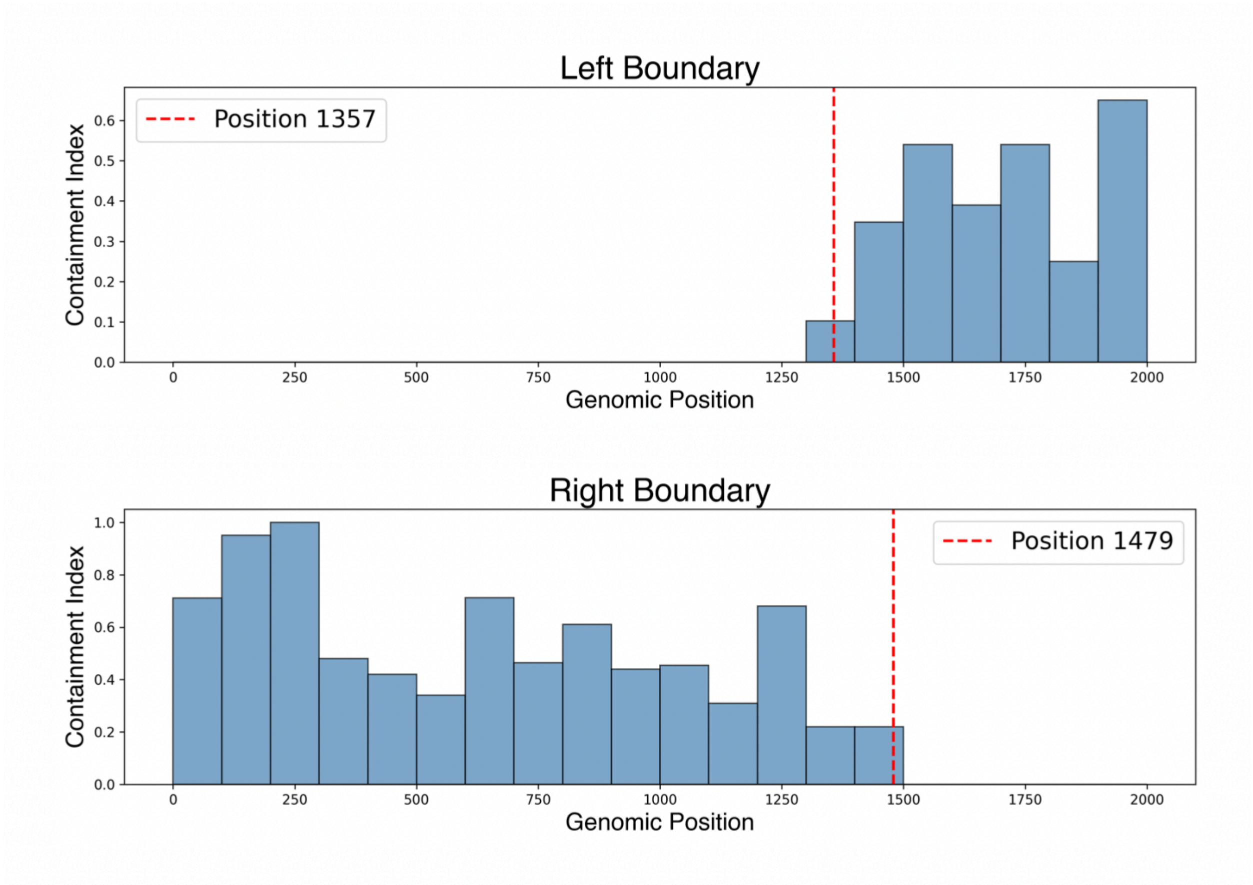


**Supplementary Figure 1:** Visualization of boundary refinement for an HSat1A array (HG002 chr13_MATERNAL:14534346-15946973). *K*-mers within the array are separated into two flanking regions (*L* and *R)*, with the remaining *k*-mers comprising the core set. *L* and *R* are further subdivided into windows (*w* = 100 bp), and each window is assigned a containment score based on the fraction of its *k*-mers present in the core set. The array boundaries are then refined by identifying the left-most window of *L* and right-most window of *R* that exceed a user-defined containment threshold, and narrowing the start and end positions to the last core *k*-mer match within those windows.

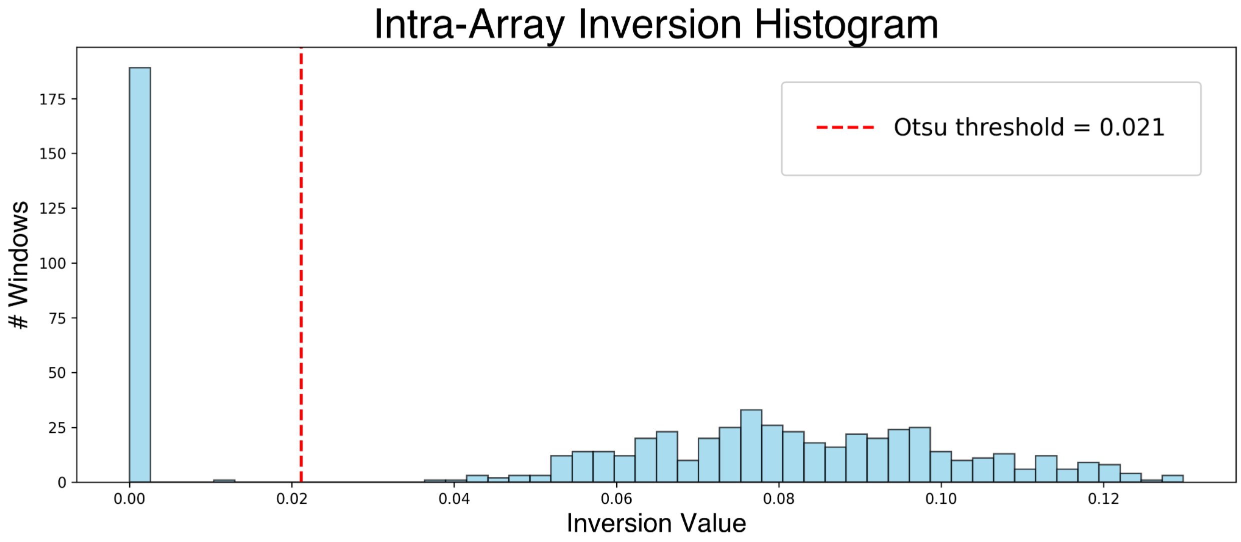


**Supplementary Figure 2:** Histogram of windowed (*w* = 1000 bp) intra-array inversion values for an HSat3 array (HG002 chr17_MATERNAL:21,673,043-22,329,332). Windows on the left hand side of Otsu's threshold are considered forward-oriented, while those on the right-hand side are considered reverse-oriented.

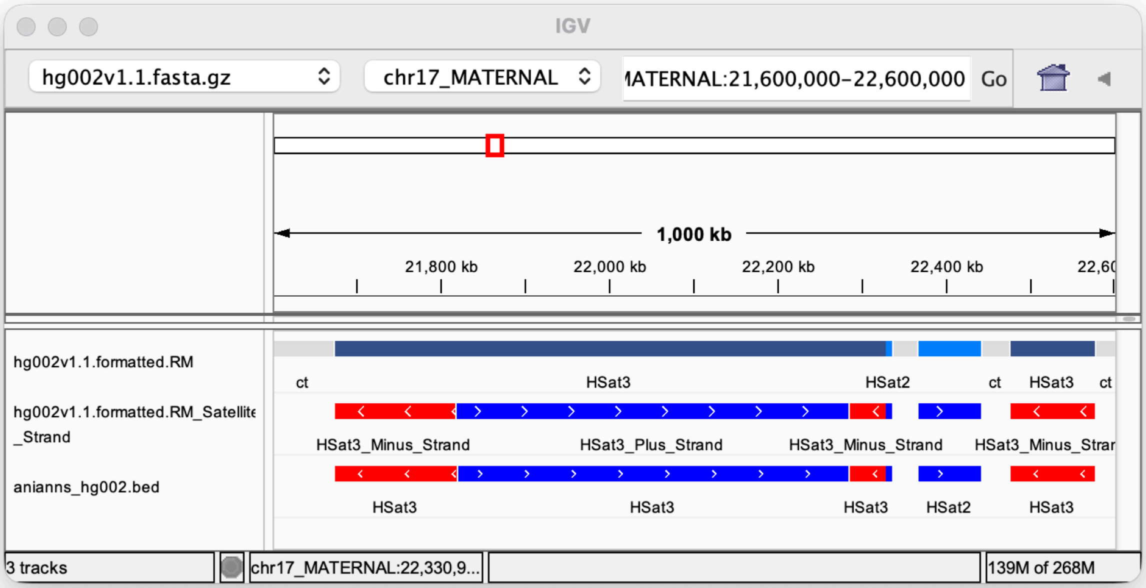


**Supplementary Figure 3:** IGV [Robinson et al., PMID: 21221095] screenshot of HG002 chr17_MATERNAL:21,600,600-22,600,000, with CenSat non-stranded (top), CenSat stranded (middle), and AniAnn's stranded (bottom) annotation tracks. Both stranded annotation tracks agree on two inversion points within the large HSat3 array.

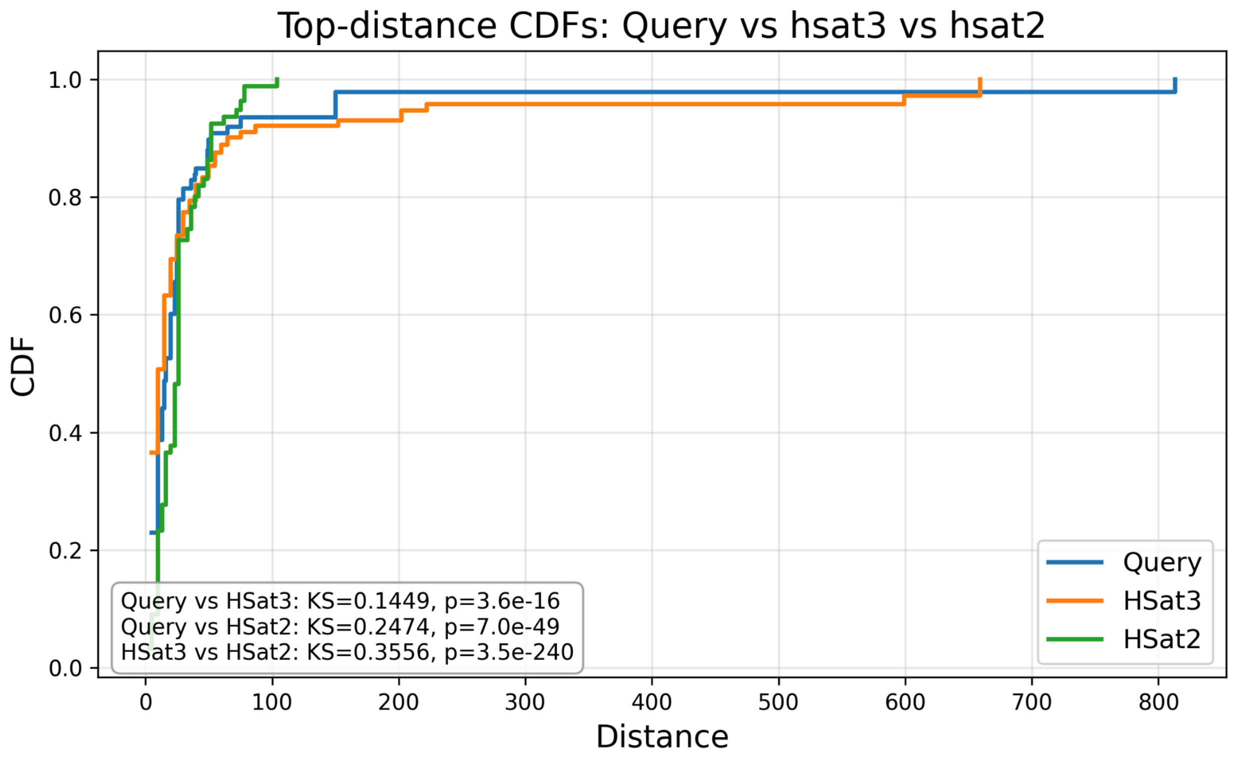


**Supplementary Figure 4:** Weighted empirical CDFs of inter-*k*-mer distance distributions for a query array and two reference arrays (HSat2 and HSat3). Distances are weighted by occurrence counts prior to CDF construction, and pairwise Kolmogorov–Smirnov (KS) tests are used to quantify distributional similarity. KS statistics and corresponding p-values are shown within the figure (bottom left).

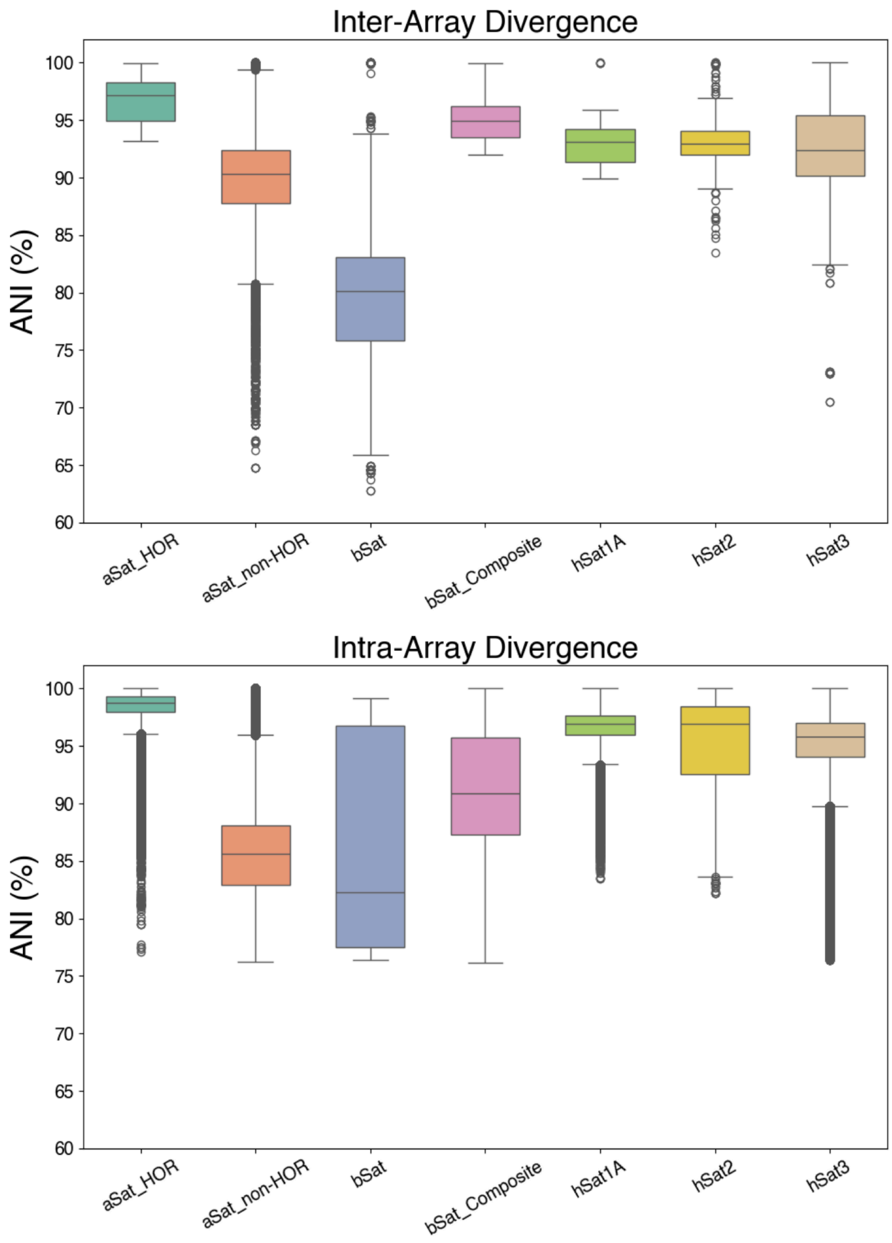


**Supplementary Figure 5:** Boxplot of inter-array and intra-array divergence values for satellite arrays within HG002. Arrays were based on their classification in theHG002 CenSat track. αSat was segregated into active HOR forming and non-HOR forming (active_hor and mon in the CenSat track, respectively), while βSat was segregated into arrays forming a composite with LSAU and those not (bSat(BSR_Beta,COMP*) and bSat(BSR_Beta), respectively). Inter-array divergence was calculated by computing the Mash distance using the full (unsketched) set of *k*-mers between each pairwise combination of arrays on each chromosome. Intra-array divergence was computed by running ModDotPlot on each array and summing the ANI values within each matrix not on the diagonal. The line inside each boxplot represents the median, while the box represents the interquartile range (IQR). Whiskers are defined at positions ± 1.5 x IQR. Outliers are shown as gray circles outside the whisker range.

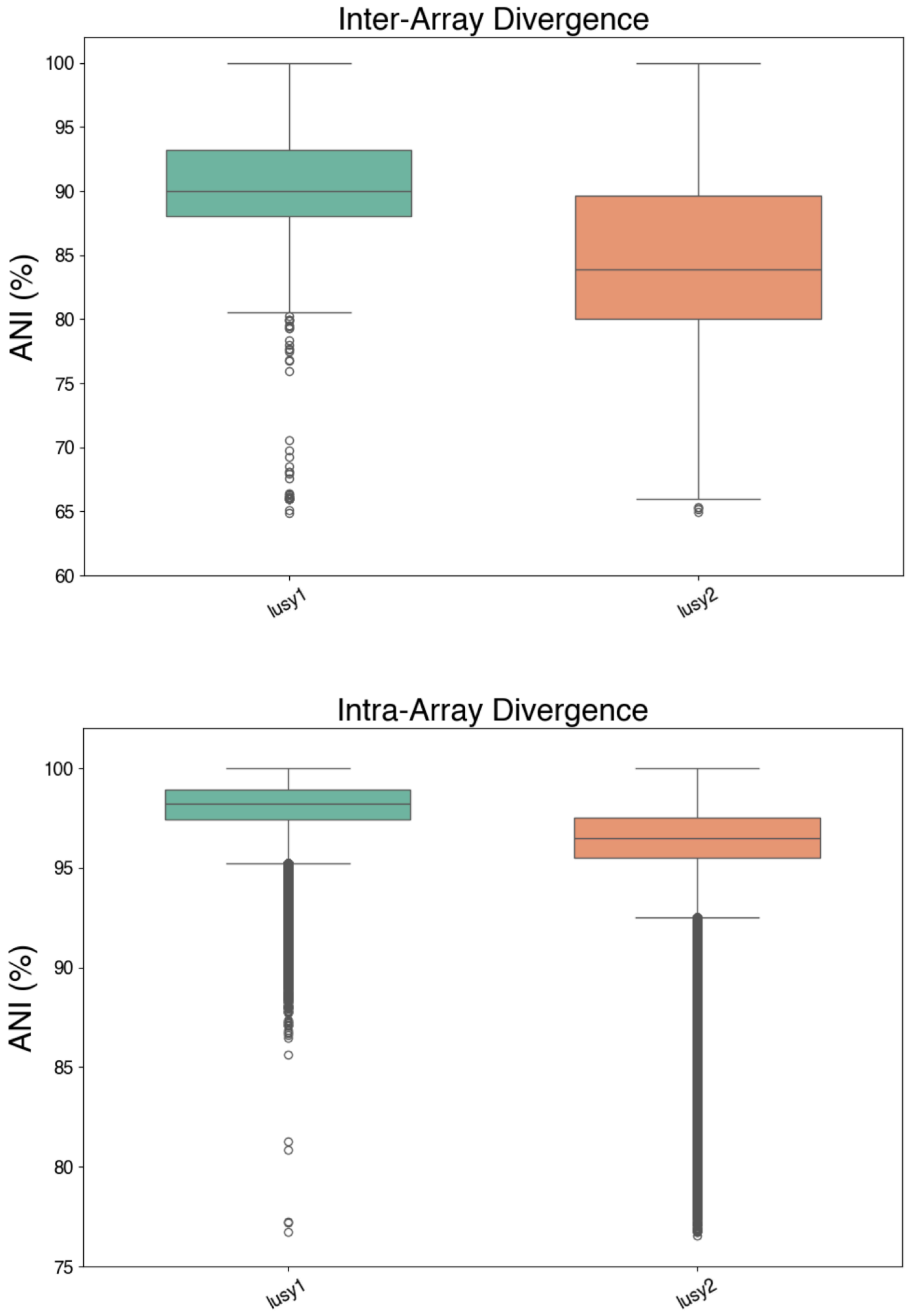


**Supplementary Figure 6:** Boxplot of inter-array and intra-array divergence values for *Lusy1* and *Lusy2* within IpLuzLuz (common woodrush). Arrays were based on their classification in the HG002 CenSat track. Inter-array divergence was calculated by computing the Mash distance using the full (unsketched) set of *k*-mers between each pairwise combination of arrays on each chromosome. Intra-array divergence was computed by running ModDotPlot on each array and summing the ANI values within each matrix not on the diagonal. The line inside each boxplot represents the median, while the box represents the interquartile range (IQR). Whiskers are defined at positions ± 1.5 x IQR. Outliers are shown as gray circles outside the whisker range.
