## Supplementary Tables for "AniAnn’s: alignment-free annotation of tandem repeat arrays using fast average nucleotide identity estimates"

| Analysis | Training Data Location | K-mer db size (M) |
|---|---|---|
| HG002 Whole Genome | https://github.com/hloucks/CenSatData/blob/main/CHM13/chm13v2.0.cenSat.v2.0.bed | 31.2 |
| mGorGor1 Whole Genome | https://github.com/hloucks/CenSatData/blob/main/T2TPrimates/PanPan/PanPan.cenSatv2.0.bed | 51.3 |
| mLuzLuz1 Whole Genome | https://zenodo.org/records/14007621/files/Luzula_sylvatica_asm.hic.chr.FINAL.TideCluster_tandem_repeats.gff3?download=1 | 19.8 |

**Supplementary Table 1:** Training data locations and k-mer db sizes for AniAnn's.

| Software Availability | Version | Command line |
|---|---|---|
| AniAnns<br>https://github.com/marbl/aniannns | v0.5.0 | `anianns annotate -f chr21_*.fa --identity 86 --mask` |
| | | `anianns annotate -f chr21_*.fa --identity 92 --mask` |
| Longdust<br>https://github.com/lh3/longdust | v1.1 | `longdust chr21_*.fa` |
| | | `longdust chr21_*.fa -t 0.3` |
| TRF<br>https://github.com/Benson-Genomics-Lab/TRF | v4.09.1 | `trf chr21_*.fa 2 7 7 80 10 50 500` |
| | | `trf chr21_*.fa 2 7 7 80 10 50 2000` |
| TRASH<br>https://github.com/vlothec/TRASH_2 | v2.0.0 | `Rscript TRASH.r --min_rep_size 5 --cores_no 32` |
| | | `Rscript TRASH.r --max_rep_size 2000 --min_rep_size 5 --cores_no 32` |
| Lastz<br>https://github.com/lastz/lastz | v1.04.52 | `lastz \`<br>` chr21_*mat_masked.fa \`<br>` chr21_*pat_masked.fa \`<br>` --notransition \`<br>` --step=1 \`<br>` --seed=match14 \`<br>` --nogapped \`<br>` --hspthresh=top95% \`<br>` --nochain \`<br>` --format=maf` |

**Supplementary Table 2:** Command line parameters used for each tool. A custom Python script was used to convert bed files into masked fasta files, and is available at github.com/alexsweeten/satelliteannotation

| Chr | Superfamily (SF) | Maternal most frequent HOR | Paternal most frequent HOR |
|---|---|---|---|
| 1 | 1 | 2 | 2 |
| 2 | 2 | 4 | 4 |
| 3 | 1 | 17 | 17 |
| 4 | 2 | 19 | 19 |
| 5 | 1 | 2 | 2 |
| 6 | 1 | 18 | 18 |
| 7 | 1 | 6 | 6 |
| 8 | 2 | 11 | 11 |
| 9 | 2 | 4 | 4 |
| 10 | 1 | 6 | 6 |
| 11 | 3 | 5 | 5 |
| 12 | 1 | 8 | 8 |
| 13 | 2 | 7 | 11 |
| 14 | 2 | 8 | 8 |
| 15 | 2 | 15 | 15 |
| 16 | 1 | 10 | 10 |
| 17 | 3 | 16 | 16 |
| 18 | 2 | 12 | 12 |
| 19 | 1 | 2 | 2 |
| 20 | 2 | 16 | 16 |
| 21 | 2 | 11 | 11 |
| 22 | 2 | 8 | 8 |
| X | 3 | 12 | - |
| Y | 4 | - | 34 |

**Supplementary Table 3:** AniAnn's α-satellite higher-order repeat (HOR) and superfamily (SF) identification across the diploid HG002 (human) genome. For each chromosome, the most frequent HOR detected in the maternal and paternal haplotypes is reported, along with the corresponding α-satellite superfamily. Differences between haplotypes are highlighted in red and reflect differences in array dominance between haplotypes.

| Chr | Superfamily (SF) | Maternal most frequent HOR | Paternal most frequent HOR |
|---|---|---|---|
| 1_hsa1 | 1 | 2 | 2 |
| 2_hsa3 | 1 | 2 | 2 |
| 3_hsa4 | 1 | 2 | 2 |
| 4_hsa17x5 | 1 | 2 | 2 |
| 5_hsa6 | 1 | 2 | 2 |
| 6_hsa7 | 1 | 2 | 2 |
| 7_hsa8 | 1 | 2 | 2 |
| 8_hsa10 | 1 | 2 | 2 |
| 9_hsa11 | 1 | 2 | 2 |
| 10_hsa12 | 1 | 2 | 2 |
| 11_hsa2b | 1 | 2 | 2 |
| 12_hsa2a | 2 | 6 | 6 |
| 13_hsa9 | 2 | 2 | 2 |
| 14_hsa13 | 2 | 5 | 5 |
| 15_hsa14 | 2 | 10 | 10 |
| 16_hsa15 | 2 | 3 | 17 |
| 17_hsa18 | 2 | 4 | 4 |
| 18_hsa16 | 1 | 2 | 2 |
| 19_hsa5x17 | 2 | 8 | 8 |
| 20_hsa19 | 1 | 2 | 2 |
| 21_hsa20 | 1 | 2 | 2 |
| 22_hsa21 | 2 | 2 | 2 |
| 23_hsa22 | 2 | 25 | 12 |
| X | 1 | 4 | - |
| Y | 1 | - | 18 |

**Supplementary Table 4:** AniAnn's α-satellite higher-order repeat (HOR) and superfamily (SF) identification across the diploid mGorGor1 (gorilla) genome. For each chromosome, the most frequent HOR detected in the maternal and paternal haplotypes is reported, along with the corresponding α-satellite superfamily. Differences between haplotypes are highlighted in red and reflect differences in array dominance between haplotypes.
